# DIANA: An integrated pipeline for analysis of long-read whole-genome sequencing data for molecular neuropathology

**DOI:** 10.64898/2026.03.25.714119

**Authors:** Christian Domilongo Bope, Henning Leske, Richard Mark Nagymihaly, Einar Osland Vik-Mo, Skarphéðinn Halldórsson

## Abstract

**Summary:** Central nervous system (CNS) tumor diagnosis requires comprehensive genomic profiling including DNA-methylation classification, copy-number variants (CNV), gene fusion analysis, small variant detection and *MGMT* promoter methylation status. Long-read sequencing platforms such as nanopore sequencing by Oxford Nanopore Technologies and SMRTseq by PacBio can capture all these in a single assay, but integrating diverse analytical tools to leverage the advantages of long-read sequencing remains complex. We present DIANA (<u>D</u>iagnostic <u>I</u>ntegrated <u>A</u>nalytics of <u>N</u>eoplastic <u>A</u>lterations), a pipeline providing fully automated end-to-end processing of long-read whole-genome sequencing data from aligned sequence reads. DIANA produces a human readable report that combines methylation classification with prioritized genetic variants to support CNS tumor diagnostics and clinical decision-making.

**Availability and implementation:** DIANA is an open-source Nextflow pipeline, freely available through Docker or Apptainer/Singularity technologies. The source code, comprehensive documentation, and installation protocols are available on GitHub: https://github.com/VilhelmMagnusLab/DIANA.git.

**Supplementary information:** Supplementary data are available at Bioinformatics online.

## INTRODUCTION

Accurate diagnosis of central nervous system (CNS) tumors requires comprehensive genetic analysis, including the detection of single nucleotide variants (SNVs), copy number variations (CNVs), gene fusions and epigenetic modifications (Louis, et al., 2021). Recently, DNA methylation profiling has emerged as a powerful tool and is now integral to the classification of some CNS tumors according to the World Health Organization (WHO) 2021 guidelines (Capper, et al., 2018). To fully capture all necessary alterations, current diagnostic workflows rely on separate platforms including next-generation sequencing, methylation arrays, pyro-sequencing, and fluorescent in situ hybridization (Bertero, et al., 2024). This creates significant logistical and financial burdens that can limit both their implementation and their routine use in clinical practice.

Long-read whole-genome sequencing captures DNA methylation, structural variants, small variants and copy number alterations simultaneously within a single assay (O’Neill, et al., 2024). Despite these advantages, integrating diverse analytical tools and interpretation of the results remains challenging.

To address these limitations, we developed DIANA, a Nextflow-based pipeline that provides end-to-end automation of long-read WGS data from CNS tumors. To our knowledge, this is the first publicly available end-to-end pipeline for tumor diagnostics using long-read WGS data. The pipeline integrates DNA methylation-based tumor classification, O^6^-methylguanine-DNA methyltransferase (*MGMT*) promoter methylation quantification, and detection and annotation of structural variants, copy number variations, and single nucleotide variants into a unified workflow. Each analytical module produces structured raw result files covering all variant categories, methylation calls, and copy number profiles. These raw outputs are subsequently aggregated, annotated and filtered to generate a concise genetic report that highlights the key diagnostic findings most relevant to patient management. All data remain fully accessible, enabling clinicians and researchers to perform independent verification or custom downstream analyses as required. Full containerization ensures reproducibility and portability across diverse computing environments, from local workstations to high-performance computing (HPC) clusters.

## DIANA workflow

DIANA pipeline is implemented in Nextflow (Langer, et al., 2025) (ver. 25.04.6) with DSL 2 enabled, integrating custom Python, R, and Bash scripts into a unified, modular workflow that enables scalable execution across local workstations and HPC clusters. The pipeline consists of four main modules that can be executed independently or sequentially as illustrated in Figure 1. The input consists of single or multiple BAM files generated by long-read sequencing from either Oxford Nanopore Technologies (ONT) or PacBio. All outputs are consolidated into a comprehensive PDF report generated with R language and R Markdown templates.

**Figure 1.**
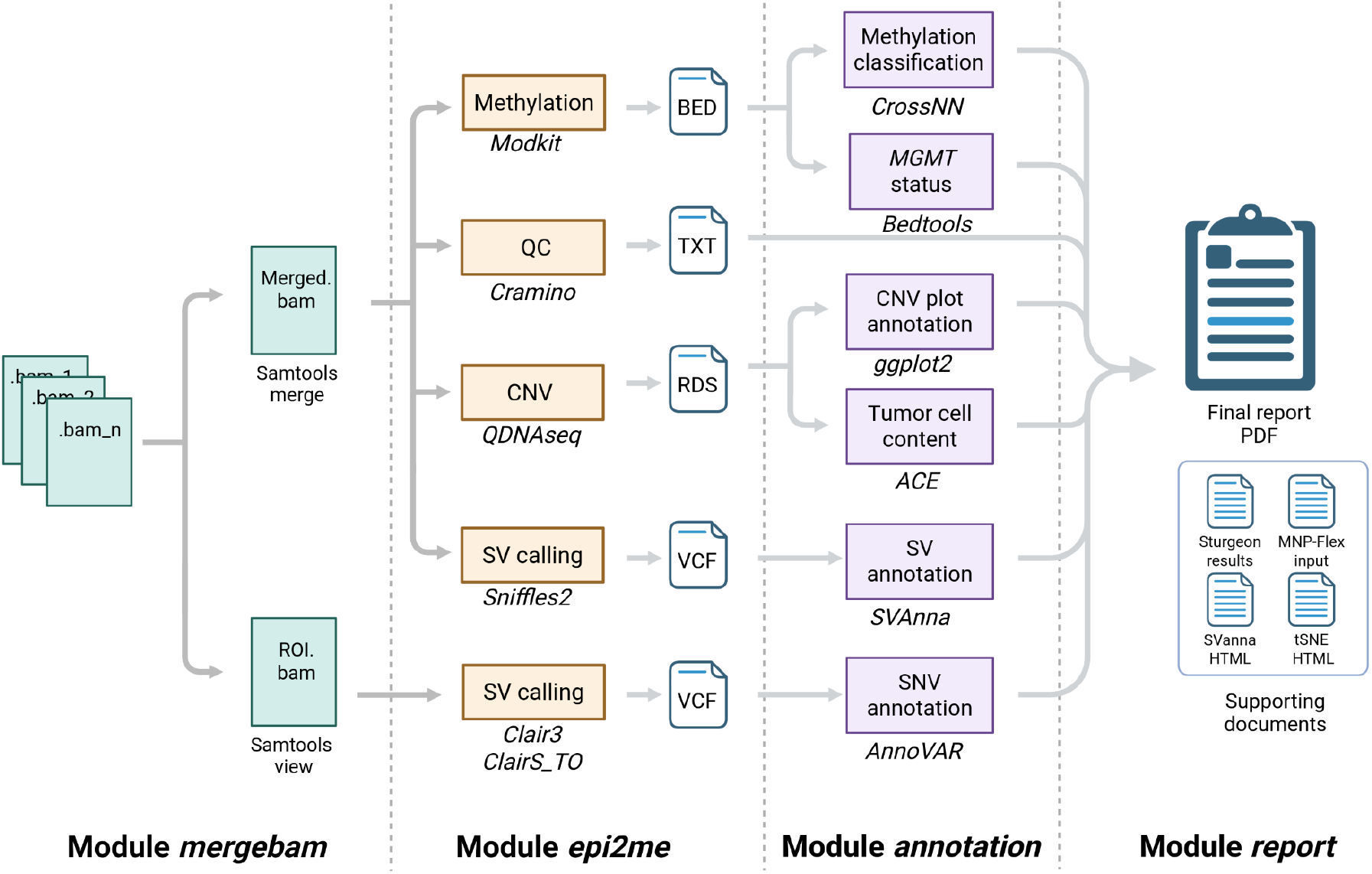
Overview of the DIANA workflow. The pipeline is organized into four distinct modules: *mergebam* for data preprocessing and region-of-interest (ROI) extraction; *epi2me* for primary genomic and epigenetic analysis, including methylation profiling (*Modkit*), structural variant calling (*Sniffles2*), and copy number variation (*QDNAseq*); *annotation* for biological interpretation and diagnostic relevance and *report* for the automated synthesis of results into a final PDF report and supporting documentation.

### First module, *mergebam*

The MinKNOW software (ONT) can be configured to write BAM files at regular intervals throughout the run (e.g., hourly) rather than producing a single file upon sequencing completion. This is the recommended approach, as it protects against data loss in the event of an unexpected interruption before the run ends. To avoid any risk of sample mix-up, the *mergebam* module uses a combination of sample identifier and flow cell identifier to unambiguously assign each BAM file to the correct sample. All files that share the same sample and flow cell identifiers are merged into a single merged and indexed BAM using *samtools merge* and *samtools index* (Danecek, et al., 2021). A second output BAM is generated by extracting only the genomic regions of interest (ROI) defined in a user-supplied BED file of clinically relevant genomic regions for SNV calling.

### Second module, *epi2me*

This module performs methylation tag extraction, structural variant calling, copy number analysis and small variant calling in parallel. Genome-wide CpG methylation quantification is performed using *modkit* (https://github.com/nanoporetech/modkit) on the whole-genome merged BAM file. Modkit interrogates the base modification tags embedded in the BAM file to produce a bedmethyl file per-CpG quantifying levels of 5-methylcytosine (*5mC*) and 5-hydroxymethylcytosine (*5hmC*) throughout the genome. Structural variant (SV) detection is carried out with *Sniffles2* (Smolka, et al., 2024) to identify insertions, deletions, inversions, and translocations and outputs a compressed VCF file. Genome-wide copy number profiling is performed using *QDNAseq* (Scheinin, et al., 2014), with a default binsize set to 50 kbp. *QDNAseq* applies GC-content and mappability correction to call copy number segments; the resulting segment BED file, bin-level BED file, and RDS object are stored as the primary raw input for the downstream analysis. Small variant calling (SNV and indel) is performed on the targeted ROI BAM file using *Clair3* (Zheng, et al., 2022) for germline variants and C*lairS-*TO (Chen, et al., 2025) for somatic variant detection. This generates compressed VCF files for downstream analysis. Sequencing quality metrics (read quality, coverage, read-length distribution) are computed using *cramino* (De Coster and Rademakers, 2023).

### Third module, *annotation*

This module takes the outputs of the *epi2me* module and performs a set of deeper interpretation and annotation steps. The following analysis is performed: (1) *MGMT* promoter methylation is quantified by extracting CpG sites overlapping the CpG island in the *MGMT* promoter, computing the mean methylation and providing a methylated versus unmethylated status (Halldorsson, et al., 2024). Results are saved as a CSV file. (2) Methylation-based classification is performed by two independent neural network classifiers: *CrossNN* (Yuan, et al., 2025) and *Sturgeon* (Vermeulen, et al., 2023). By default, both classifiers use the reference data provided by Capper *et al*. (Capper, et al., 2018) to assign a 2021 WHO tumour class with probability scores. In addition, *CrossNN* can be set to use the Pan-cancer reference data for broad tumor type classification (using the *--pancan* flag). A *UMAP* plot is also generated placing the sample context with the reference data. (3) Structural variant annotation is carried out with *Svanna* (Danis, et al., 2022), which cross-references the *Sniffles2* VCF against Human Phenotype Ontology (HPO) based gene prioritisation to highlight clinically relevant rearrangements. Gene fusion events with exon-level breakpoint coordinates are extracted and specifically annotated. (4) An annotated CNV plot is generated from the *QDNAseq* segment and bin outputs which places the corrected *log2ratio* of each bin and segment onto chromosomal coordinates using ggplot2 in R. Genes or gene-pairs of interest are annotated according to a user provided BED file. Chromosomal segments with gains or losses are highlighted. (5) The RDS file generated by *QDNAseq* is used as input for tumour cell content estimation using *ACE* (Poell, et al., 2019). (6) The small variant VCF files generated by *Clair3* and *ClairS_TO* are merged, filtered, and annotated against *ClinVar* (Landrum, et al., 2025) and *COSMIC* databases (Tate, et al., 2019) to produce a single ranked variant table. Integrated Genomics Viewer (*IGV)* (Thorvaldsdottir, et al., 2013) coverage snapshots are generated for key diagnostic loci (*EGFR, IDH1, IDH2, TERT* promoter).

### Fourth module, *report generation*

All outputs from the *analysis* module are consolidated into a comprehensive PDF report generated via an R Markdown template. The report is structured in two complementary sections. An Executive Summary provides a rapid clinical overview on a single page, opening with the mean sequencing coverage and the estimated tumour cell content, followed by: the methylation-based tumour *CrossNN* classification at class and class-family level with confidence scores, the genome-wide copy number profile, a filtered SNV table restricted to diagnostically relevant target genes and pathogenic *ClinVar* variants, a summary of detected fusion events, and the *MGMT* promoter methylation status. An Extended Report follows that provides full analytical detail for each domain: detailed read statistics (number of alignments, percentage of mapped reads, total read count, and mean coverage), the full tumour content estimate; *CrossNN* classification Methylation Class and Methylation Class Family bar plots for the top five classification scores; a *UMAP* dimensionality reduction plot for visual inspection of classifier positioning; all chromosome arm-level CNV alterations; a high-confidence CNV table filtered for amplifications and homozygous deletions; focused chromosome 7 (*EGFR*) and chromosome 9 (*CDKN2A/B*) profiles; overage snapshots for *IDH1* p.R132, *IDH2* p.R172, and *TERT* promoter hotspots; the full per-CpG *MGMT* methylation table (CpG sites 1–98 and 76–79); and the complete annotated SNV and structural variant tables. Alongside the main report PDF, a set of supplementary outputs is published: a *Sturgeon* methylation classification PDF, an interactive *Svanna* Structure Variant annotation HTML, an interactive *CrossNN* classification *UMAP* HTML plot, and a formatted methylation BED file for compatibility with *MNP-Flex* classification platforms.

## DIANA run-modes

DIANA supports flexible execution through multiple run modes. (i) *Smart sample monitor*: an automated monitoring utility that continuously watches sample directories for the presence of a sequencing completion marker file (*final_summary_*_*_\**.*txt*) that is written by MinKNOW when the sequencing run reaches its predefined endpoint (time or data limit reached) or is manually stopped. Upon detection, it automatically queues and launches the full pipeline from *mergebam* to R Markdown *report* for each completed sample. (ii) *–run_mode_mergebam*: runs only the BAM merging and ROI extraction step, for example when BAM preparation must be performed separately. (iii) *–run_mode_order*: triggers complete end-to-end execution of all modules in sequence (*mergebam epi2me annotation report*), processing samples from unmerged BAM files through to the final report. (iv) *–run_mode_epianalyse:* runs the *epi2me* and *annotation* modules sequentially, skipping *mergebam*; intended for cases where BAM files have already been merged. (v) Individual sub-module execution: each *epi2me* analysis type and each *annotation* step can be run independently using dedicated flags, for example *–run_mode_epi2me modkit* to rerun methylation calling only, –*run_mode_analysis mgmt* for *MGMT* quantification, and –*run_mode_analysis rmd* to regenerate the PDF report without rerunning the upstream analyses. An example report generated from a glioblastoma genome with nanopore whole-genome sequencing is found within the **Supplementary Document 1**.

## DISCUSSION

DIANA provides a reproducible and standardized framework that ensures that all diagnostically relevant dimensions of CNS tumor characterization including DNA methylation classification, *MGMT* methylation status of the promoter region, copy number alterations, structural variants and single nucleotide variants are systematically captured, annotated, and presented. The open-source, containerised design ensures that results are fully reproducible across computing environments. The modular architecture further allows individual analytical steps to be rerun in isolation when needed, without reprocessing the entire dataset, reducing both compute time and resource consumption. The complete pipeline modules (*mergebam, epi2me*, and *annotation*) for a sample with 28.78× coverage were completed in approximately 82 minutes of wall-clock time on a single workstation using parallel processing (Supplementary Fig. S1). Resource usage varied substantially across processes: the rate-limiting steps were BAM merging (19 minutes, peak RAM 1.3 GB) in the *mergebam* module and somatic SNV calling with *ClairS-TO* (59 minutes, peak RAM 28.6 GB) in the *epi2me* module. The annotation module was notably lightweight, completing in under 4 minutes for all processes. The longest individual process was the generation of *UMAP* methylation classifier visualization (1 minute 40 seconds, peak RAM 37.9 GB). Based on these benchmarks, the minimum recommended system configuration to run the full DIANA pipeline is 32 GB of RAM, 8 CPU cores, and approximately 500 GB of available disk space for intermediate files. All processes completed successfully with no out-of-memory failures, and the annotation module ran sequentially after *epi2me*, contributing less than 5% of total runtime.

Accessibility was a central design consideration: DIANA runs on standard local workstations as well as HPC clusters, requires no prior bioinformatics expertise to operate, and is freely available under MIT open licence with fully automated setup scripts.

## ACKNOWLEDGEMENTS

We thank Dr. Philipp Euskirchen and Dr. Areeba Patel for insightful discussions on methylation classifiers and analytic software, and Dr. Marius Lund Iversen for help with variant prioritization. The authors declare the use of Claude (Anthropic, Claude Sonnet 4.6) as an AI-assisted tool for code debugging during the development of the DIANA pipeline. All AI-generated contributions were reviewed and validated by the authors.

## FIGURES

**Supplementary Figure 1.**
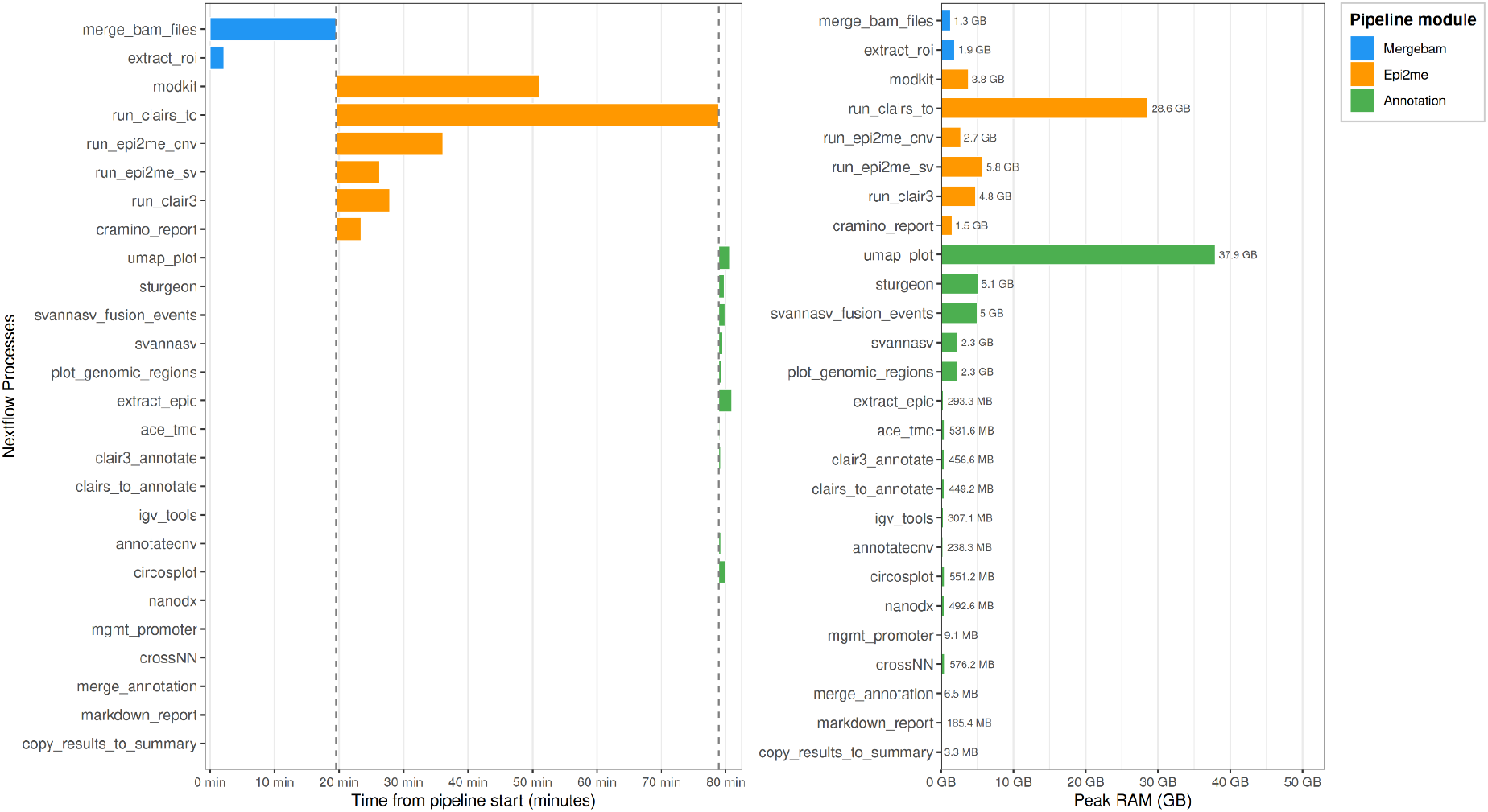
Computational resource utilization of the DIANA pipeline. **(A)** Gantt chart showing the wall-clock execution timeline of all Nextflow processes for a representative glioblastoma sample with 28.78× coverage run on a single workstation (190 GB RAM, 48 CPU cores). Processes are grouped by pipeline module: BAM preparation (***Mergebam***, blue), variant and methylation calling (***Epi2me***, orange), and report generation (***Annotation***, green). Dashed vertical lines indicate module boundaries. Within each module, processes are executed in parallel. Total pipeline runtime was approximately 82 minutes, with bam file merging (merge_bam_file, 19 minutes) *and* somatic SNV calling (run_clairs_to, 59 min) constituting the rate-limiting steps. **(B)** Peak RAM usage per process. The most memory-intensive steps were the *UMAP* plot methylation classifier visualization (umap_plot, 37.9 GB) and somatic variant calling (run_clairs_to, 28.6 GB). The majority of annotation processes required less than 1 GB of RAM.

## Executive Summary

### Sample ID: diana-008

The report is based on a mean coverage of **37.65** and the estimated tumor content in sample is **63%**.

#### CrossNN Methylation-based Classification

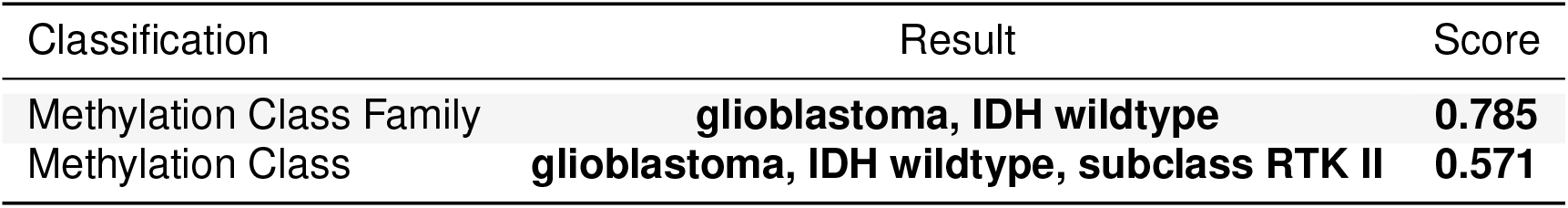

| Classification | Result | Score |
| --- | --- | --- |
| Methylation Class Family | **glioblastoma, IDH wildtype** | **0.785** |
| Methylation Class | **glioblastoma, IDH wildtype, subclass RTK II** | **0.571** |

#### Full CNV Profile

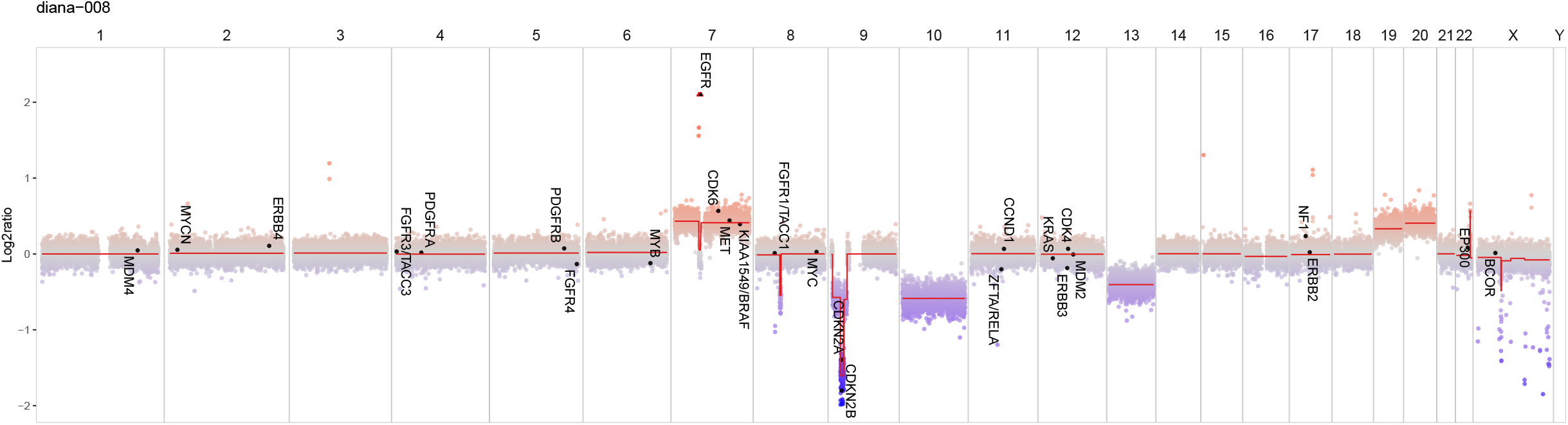

#### SNV Calling and Annotation

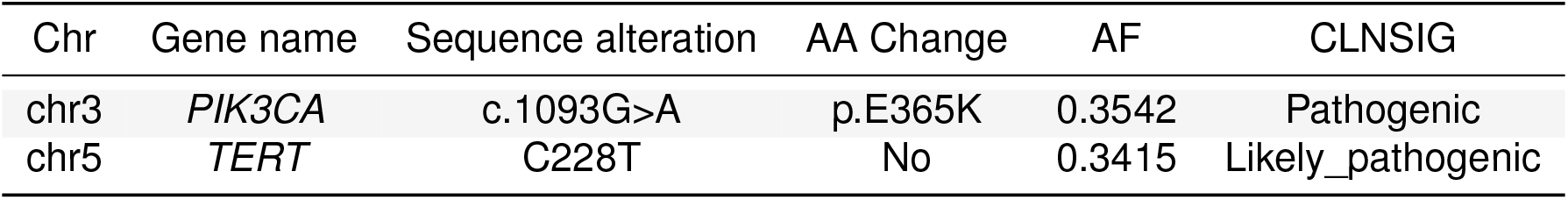

| Chr | Gene name | Sequence alteration | AA Change | AF | CLNSIG |
| --- | --- | --- | --- | --- | --- |
| chr3 | *PIK3CA* | c.1093G>A | p.E365K | 0.3542 | Pathogenic |
| chr5 | *TERT* | C228T | No | 0.3415 | Likely_pathogenic |

#### Structural Variants/Fusion Events

~~~
No fusion event break point detected in this sample.
~~~

#### *MGMT* methylation status

The mean methylation is **16.11%**. The methylation status of the *MGMT* promoter region is classified as **Unmethylated**.

## Nanopore Extended WGS Report

### Sample ID: diana-008

#### Read statistics

~~~
Number of alignments 32416596
% from total alignments 91.52
Number of reads 29750250
Mean coverage 37.65
The estimated tumor cell content for diana-008 is: 63%
~~~

#### CrossNN Methylation-based Classification

Methylation-based classification is based on **365471** CpG sites (overlap of sites covered in this sample and the model). At the methylation class (MC) level, the sample has been classified as **glioblastoma, IDH wildtype, subclass RTK II**. This prediction has a confidence score of **0.571**. At the methylation class **family** (MCF) level, the sample has been classified as **glioblastoma, IDH wildtype**. The MCF prediction has a confidence score of **0.785**.

Scores for the Top 5 entities on MC and MCF level are given below. Vertical dashed lines indicate the recommended >0.2 cut-off for classification.

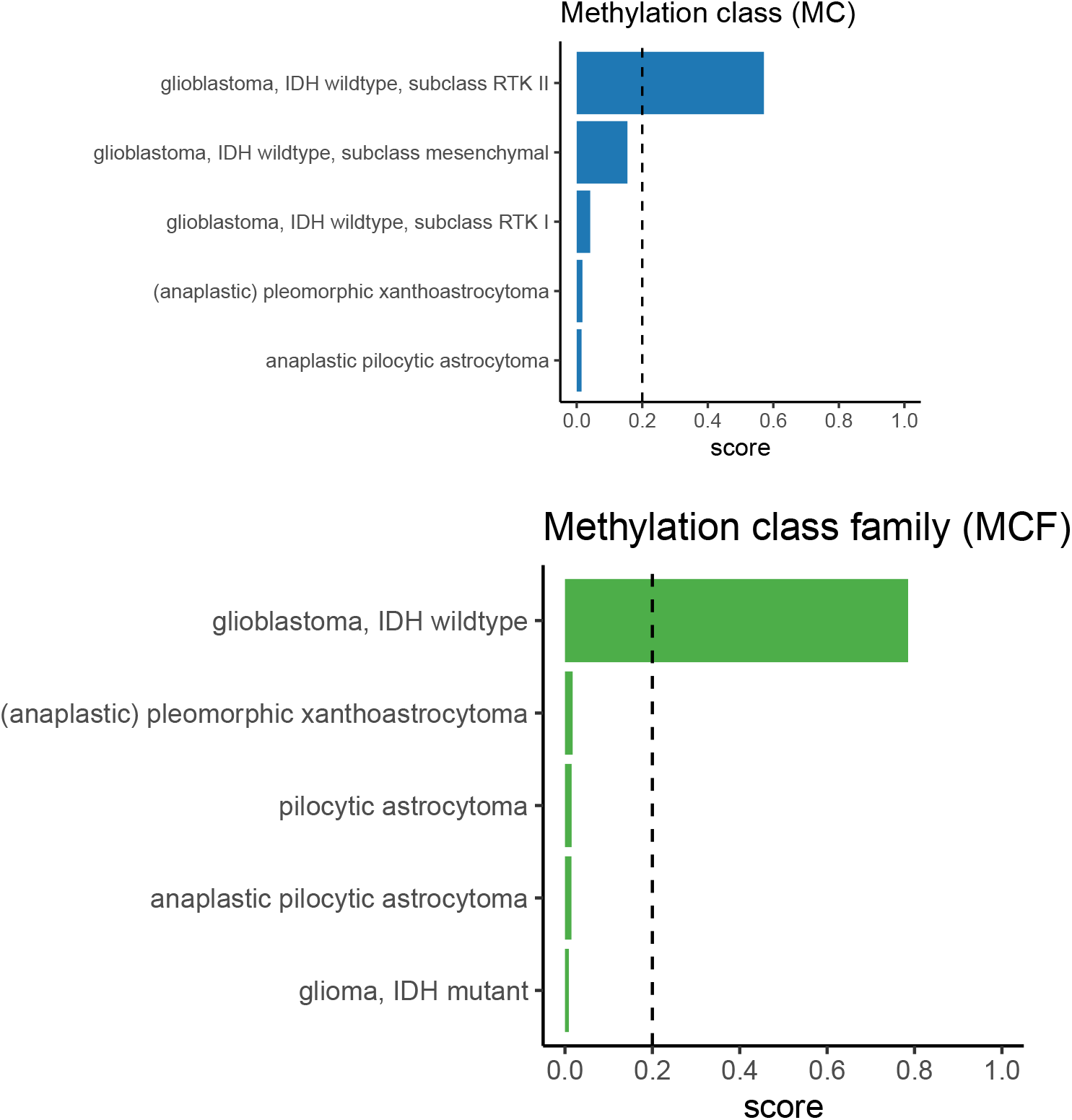

## Dimensionality Reduction Plot

The dimensionality reduction plot are only intended for visual inspection, not classification.

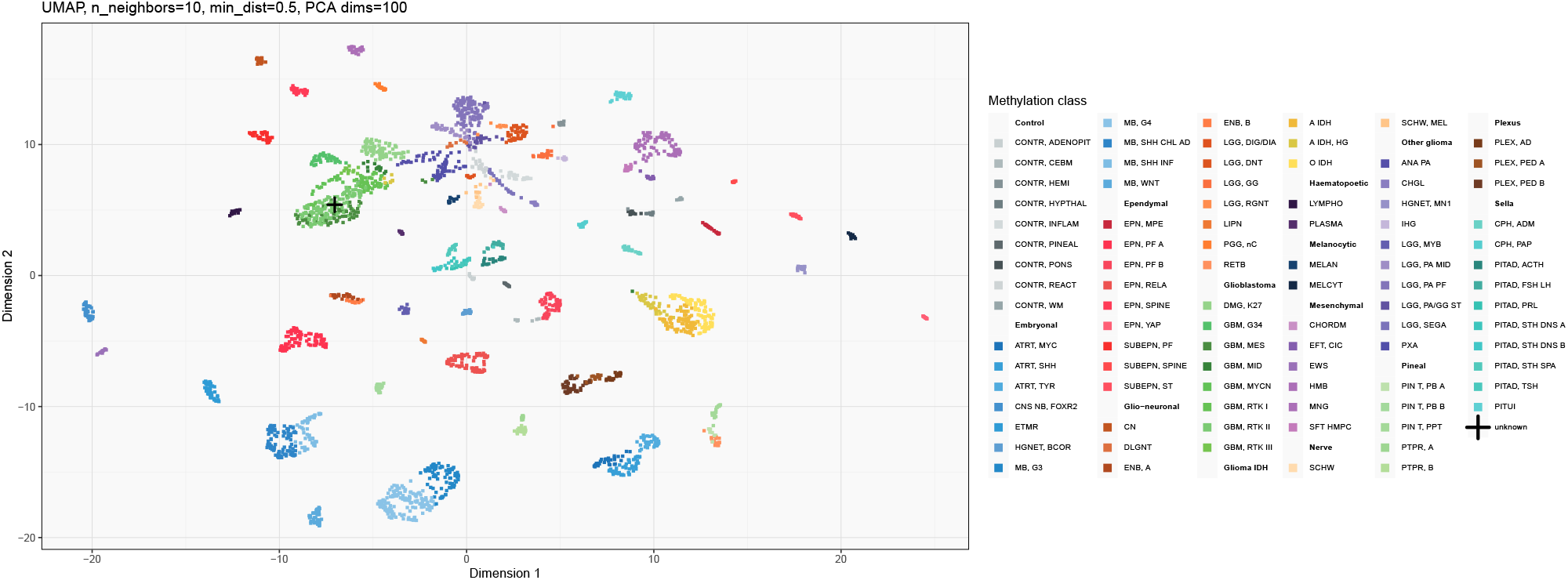

## Copy Number Variation Profile

### Full CNV Profile

~~~
Genes annotated in the full CNV profile are amplified (Gain) or deleted (Loss) based on QDNAseq results.
~~~

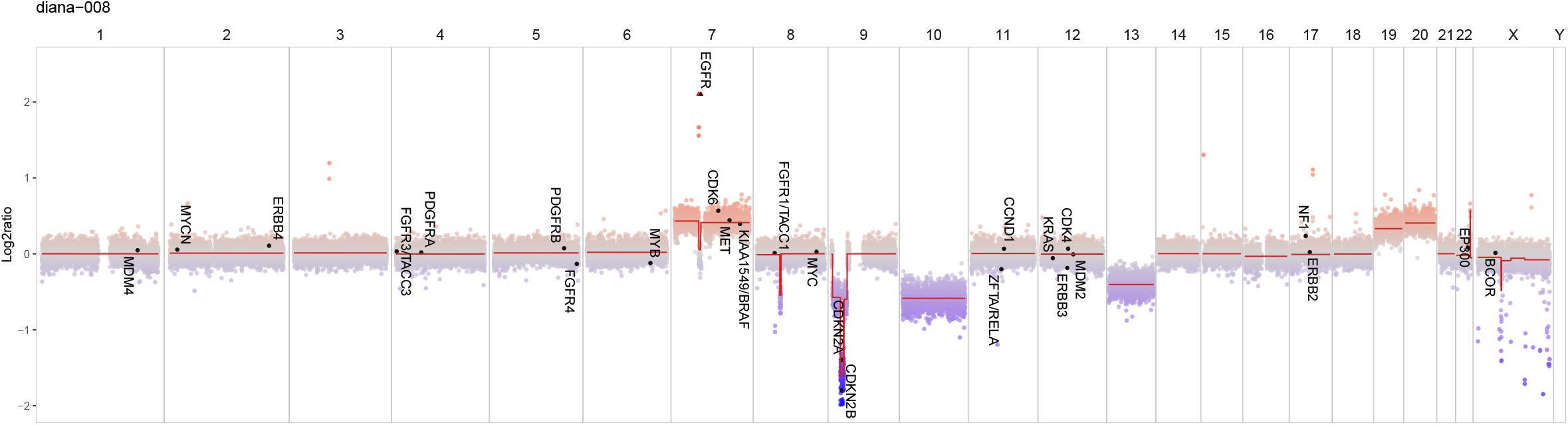

~~~
Chromosome arm-level alterations detected (>90% threshold):
~~~

~~~
- Chr7: Gain
- Chr10: Loss
- Chr13: Loss
- Chr19: Gain
- Chr20: Gain
~~~

### High-Confidence Copy Number Variation in Cancer Genes (Amplifications & Homozygous Deletions)

~~~
The table is filtered for copy number variation events with a score of 2 (amplification) or -2 (homozygous deletion) and no sex chromosomes.
~~~

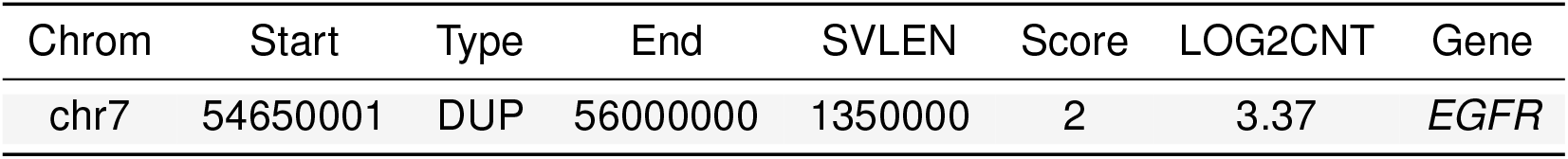

### Chromosome 7: *EGFR* Profile

Chromosome 7 CNV for visual inspection of *EGFR* annotated.

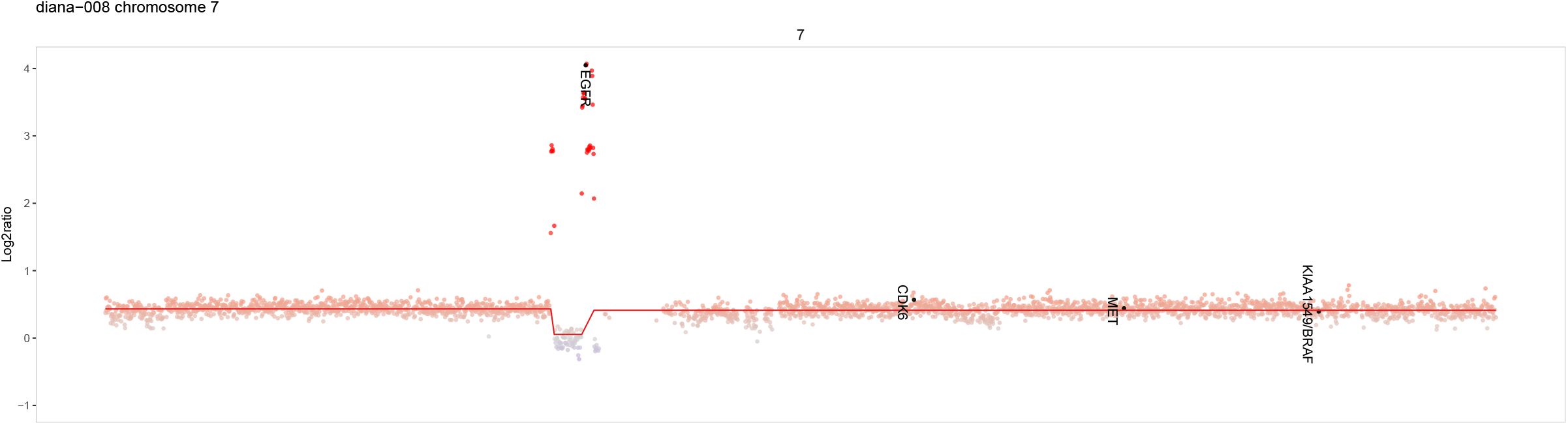

The *EGFR* copy number for the sample diana-008 is **32.00**. The vertical red line highlights exons 2-7. Deletion of exons 2-7 results in *EGFR*viii variant.

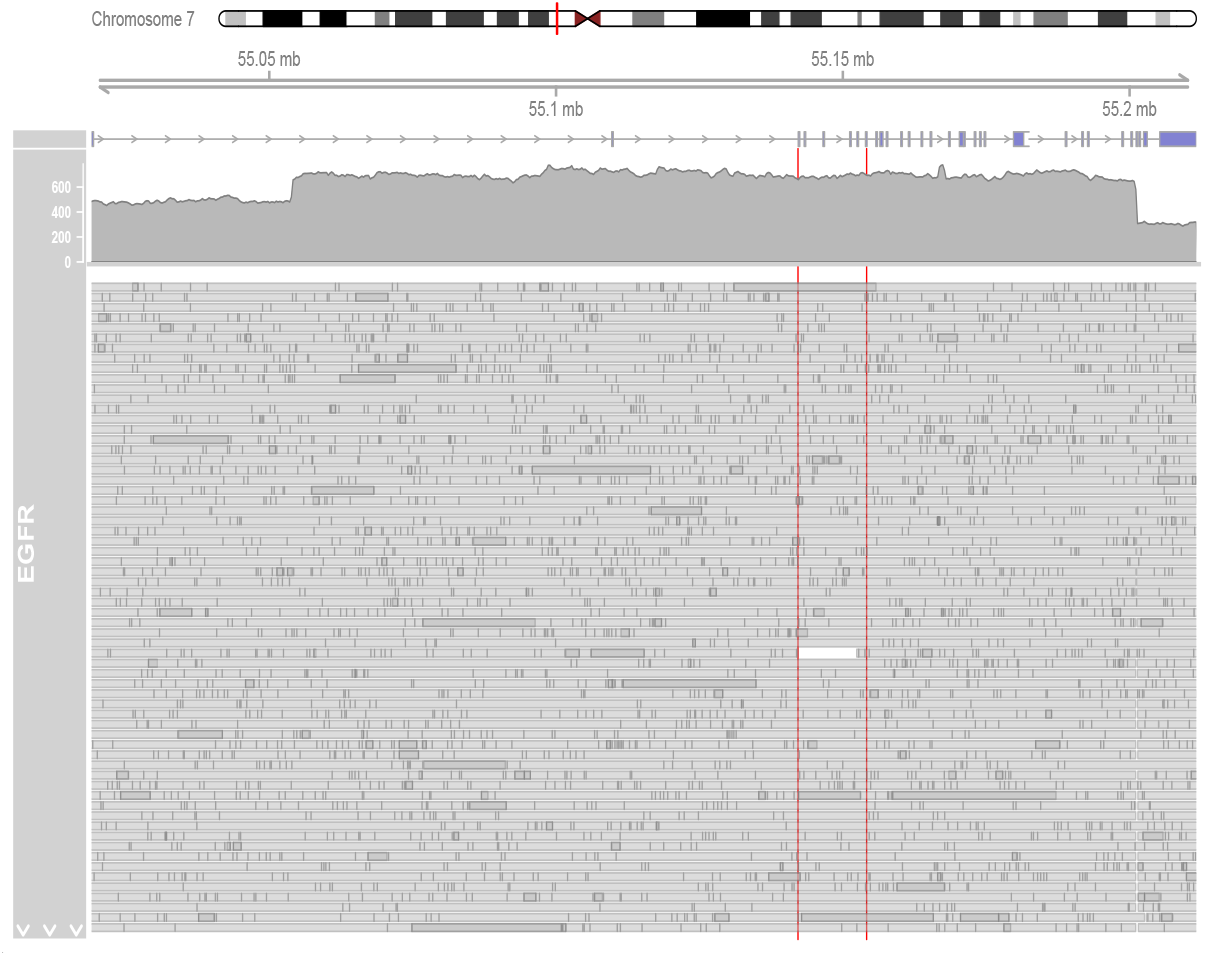

### Chromosome 9: *CDKN2A/B* Profile

Chromosome 9 CNV for visual inspection of *CDKN2A/B* annotated.

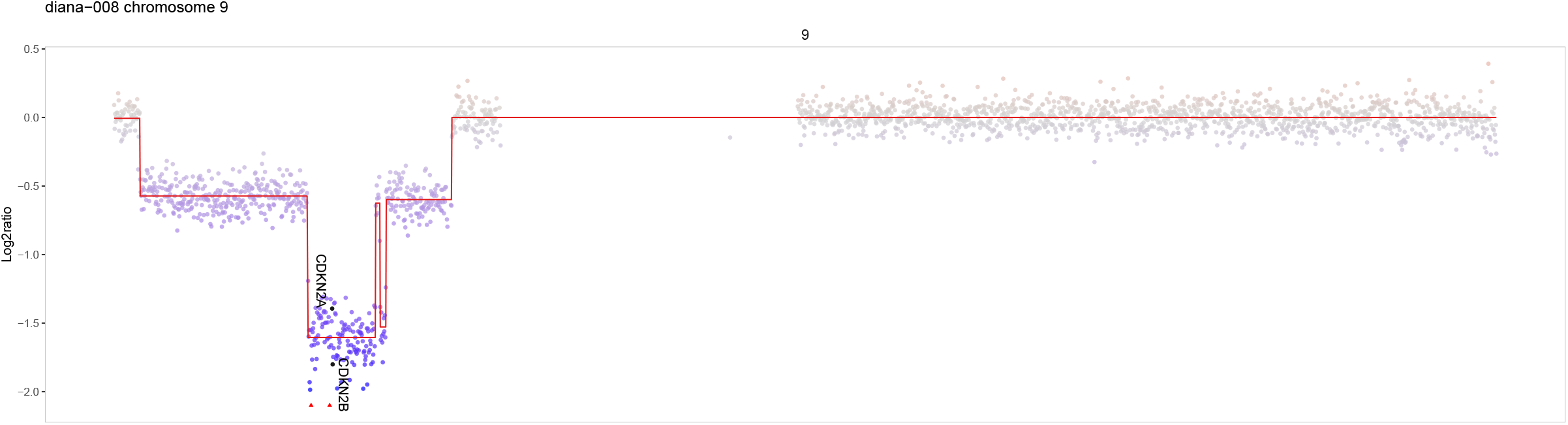

There is a homozygous deletion in chromosome 9: *CDKN2A* is deleted There is a homozygous deletion in chromosome 9:

*CDKN2B* is deleted

### *IDH1* p.R132 and *IDH2* p.R172

Sequenced coverage of *IDH1* p.R132 and *IDH2* p.R172 (red vertical line) for visual inspection.

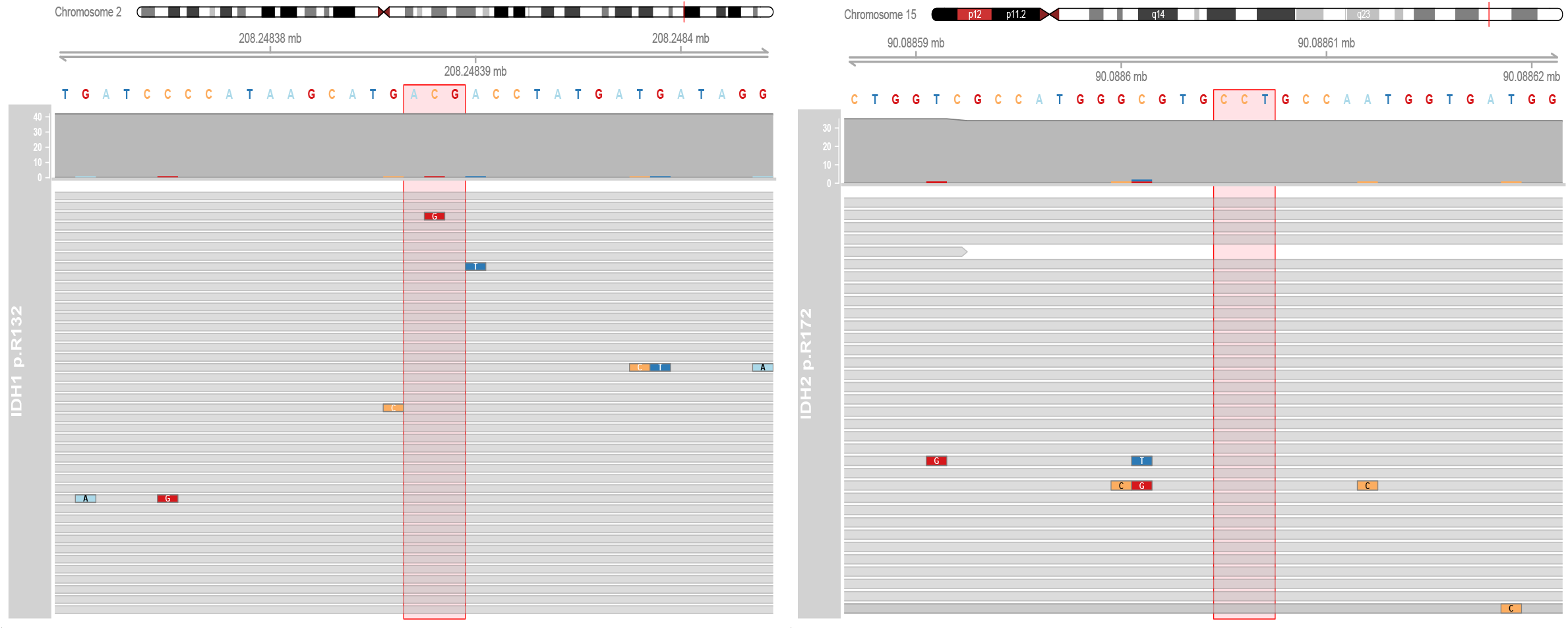

### *TERT* p

Sequenced coverage of *TERT* p C228T (left vertical red line) and C250T (right vertical red line) for visual inspection.

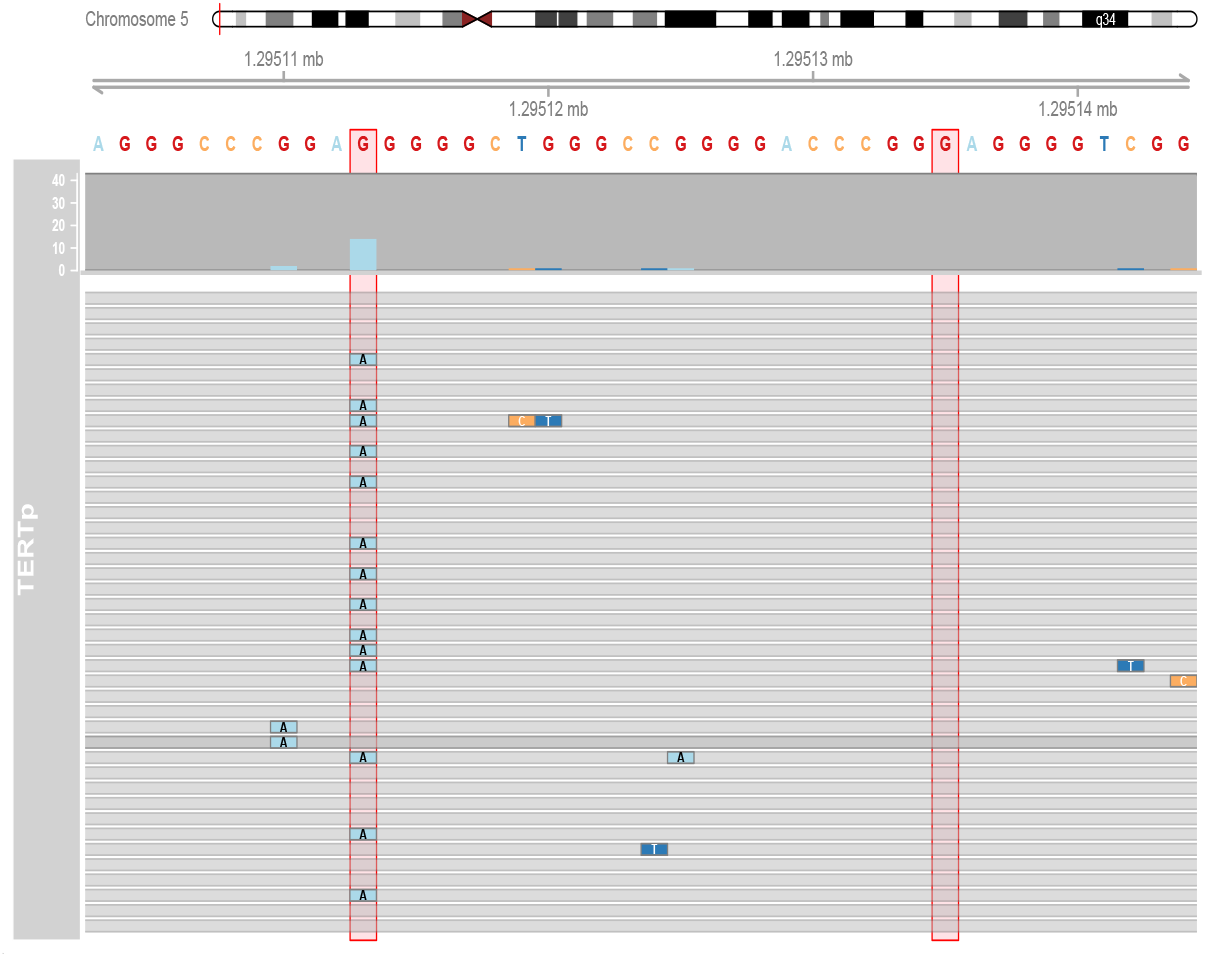

### *MGMT* methylation status

The table below summarizes methylation levels on all CpG sites in the *MGMT* promoter region (sites 1-98) and the four previously reported sites (sites 76-79) and states how the sample classifies according to both methods. Methylation above 30% in either context is considered *MGMT* methylated with high confidence. Methylation below 10% at CpGs 76-79 or below 25% at CpGs 1-98 is considered *MGMT* unmethylated with high confidence. *Grey-zone* indicates that *MGMT* methylation status can not be determined with high confidence.

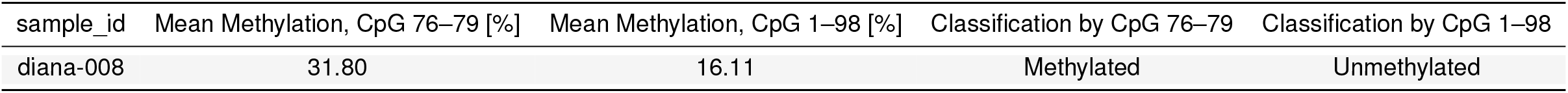

### SNV Calling and Annotation

SNVs in the select genes. Only non-synonymous exonic variants that are not known to be benign according to Clin- Var_20240611 and known pathogenic upstream variants, i.e. *TERT* p, are reported. GQ: Genotype Quality, Depth: Sequenced depth, AD: Allele Depth, GT: Allele Genotype, AF: Allele Frequency. The *GRCh38* build reference genome was used for the SNV calling and annotation. variant callers, P:Clair3 Pileup, M:Clair3 Merged and S_TO:ClairS-TO Somatic tumour-only

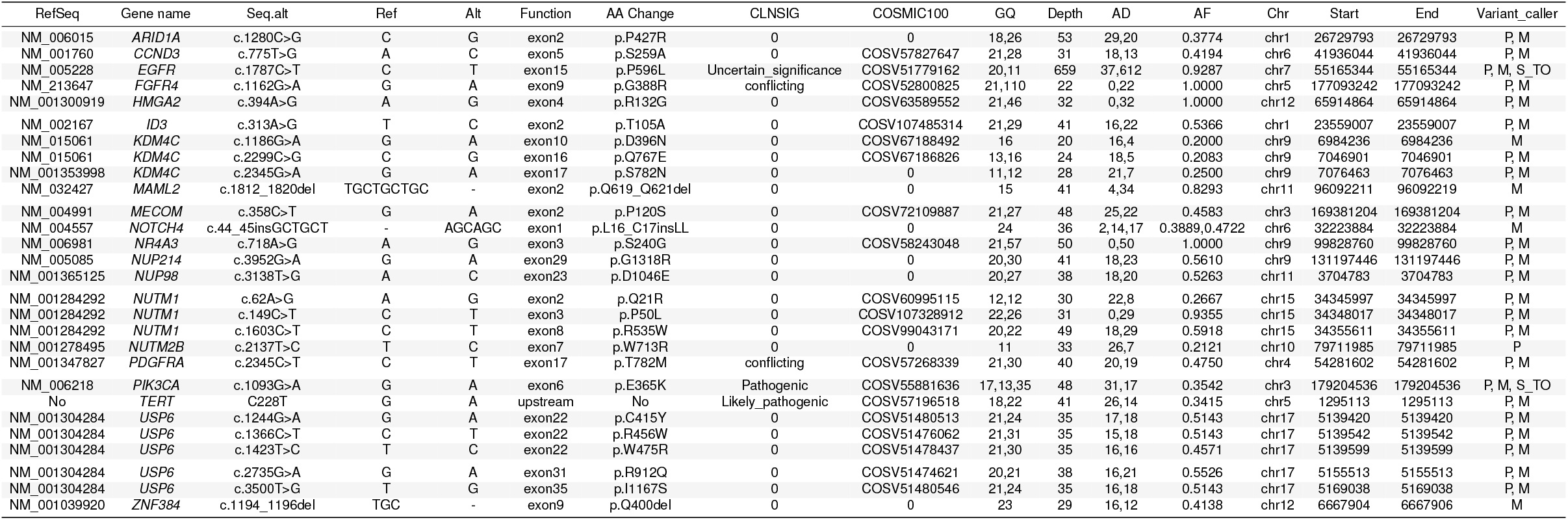

### Structural Variation/Fusion Events

~~~
No fusion event break point detected in this sample.
~~~

### Disclaimer

Diagnostic Integrated Analytics for Neoplastic Alterations pipeline (DIANA) is an investigational research tool that has not undergone full clinical validation. Any clinical use or interpretation of its results is entirely at the discretion and responsibility of the treating physician

### Report generated on 2026-03-23 08:27:48.09176 | DIANA Pipeline version: 1.0.1

